# Parental neural responsivity to infants’ visual attention: how mature brains scaffold immature brains during social interaction

**DOI:** 10.1101/295790

**Authors:** Wass S.V., Noreika V., Georgieva S., Clackson K., Brightman L., Nutbrown R., Santamaria L., Leong V.

## Abstract

Almost all attention and learning - in particular, most early learning – takes place in social settings. But little is known of how our brains support dynamic social interactions. We recorded dual-EEG from infants and parents during solo play and joint play. During solo play, fluctuations in infants’ Theta power significantly forward-predicted their subsequent attentional behaviours. But this forwards-predictiveness was lower during joint play than solo play, suggesting that infants’ endogenous neural control over attention is greater during solo play. Overall, however, infants were more attentive to the objects during joint play. To understand why, we examined how adult brain activity related to infant attention. We found that parents’ Theta power closely tracked and responded to changes in their infants’ attention. Further, instances in which parents showed greater neural responsivity were associated with longer sustained attention by infants. Our results offer new insights into how one partner influences another during social interaction.

## Introduction

Attention and learning are supported by endogenous oscillatory activity in the brain ([1–4]). The nature of these oscillations, and their relationship to behaviour, develops and changes between infancy into adulthood ([5–9]). In infants, convergent research has suggested that Theta band oscillations, which are particularly marked during early development ([10]), are associated with attentional processes. Theta band activity increases in infants during periods of anticipatory and sustained attention ([11]); in 11-month-old infants, differences in Theta band oscillations predict differential subsequent object recognition during preferential looking ([12]). Theta activity also increases in infants in social compared to non-social settings ([13]) and is particularly marked in naturalistic settings ([13]).

Although considerable previous research has investigated how brain oscillations relate to an individual’s behaviour, only a smaller body of research has investigated the neural mechanisms through which interpersonal and social factors influence behaviour ([14–16]). This is despite the fact that our brains have evolved for social living ([17]) and most of our lives – particularly early life – is spent in social settings ([18]). Understanding how social influences on attention and learning are substantiated across the brains of people engaging in social interaction, particularly during the crucial early stages of attention and learning, is an important goal for research ([19, 20]).

Previous work has shown that social factors influence infant attention and behaviour over short time-frames (seconds/minutes) and long time-frames (months/years). Over long time-frames, the children of parents who engage in more joint engagement during play show superior cognitive outcomes ([21–23]). Over short time-frames, when an infant and social partner jointly attend to the same object during naturalistic play, infant attention is increased ([24]). Recent research has contrasted two explanations for this finding: first, that social context may cause infants to be more attentive because they are more in control of their own attention behaviours. Second, that social context may offer increased opportunities for parents to scaffold their child’s attention using external attention cues – so infants are more attentive even though they are less in control of their own attention behaviours ([25]). Time-series analyses conducted to evaluate these two hypotheses provided evidence more consistent with the latter hypothesis: first, infants’ rate of change of attentiveness was faster during Joint Play than Solo Play, suggesting that internal attention factors, such as attentional inertia, may influence looking behaviour less during Joint Play ([26]). Second, adults’ attention forward-predicted infants’ subsequent attention more than *vice versa* ([25]). These results suggest that infants’ increased attentiveness during social relative to solo play may be attributable to increased attention scaffolding from parents using exogenous attention cues ([27]).

Previous research has shown that ostensive social cues such as eye gaze and vocalisations can lead to increases in inter-personal neural synchrony between infants and adults ([28]). Bidirectional Granger-causal influences between the brains of infants and adults engaged in social indirection were observed in the Theta and Alpha frequency bands, that were stronger during direct relative to indirect gaze ([28]; see also [29]; [30]). Infants vocalised more frequently during direct gaze, and individual infants who vocalised longer elicited stronger synchronisation from the adult ([28]). These findings raise the possibility that, conversely, interpersonal influences between the brains of individuals engaged in social interaction may also actively drive their partners’ attentional processes, and behaviour.

To assess this possibility, here we examined the neural and behavioural dynamics of infants’ and adults’ attention in two contexts (see Figure 1). During Joint Play, each dyad was presented consecutively with toy objects and asked to play together. During Solo Play a 40cm-high divider was placed between the infant and the parent, and two identical toys were presented concurrently to child and parent, who played separately (see Figure 1). Looking behaviour was videoed and coded *post hoc*, frame by frame, at a rate of 30Hz. Time-lagged cross-correlations were used to assess how changes in one time-series preceded or followed changes in another ([31]; cf [32, 33]) – an approach similar, but not identical, to Granger-causality ([34]). Our analyses examined whether changes in one time-series ‘forward-predicted’ changes in the other.

**Figure 1.**
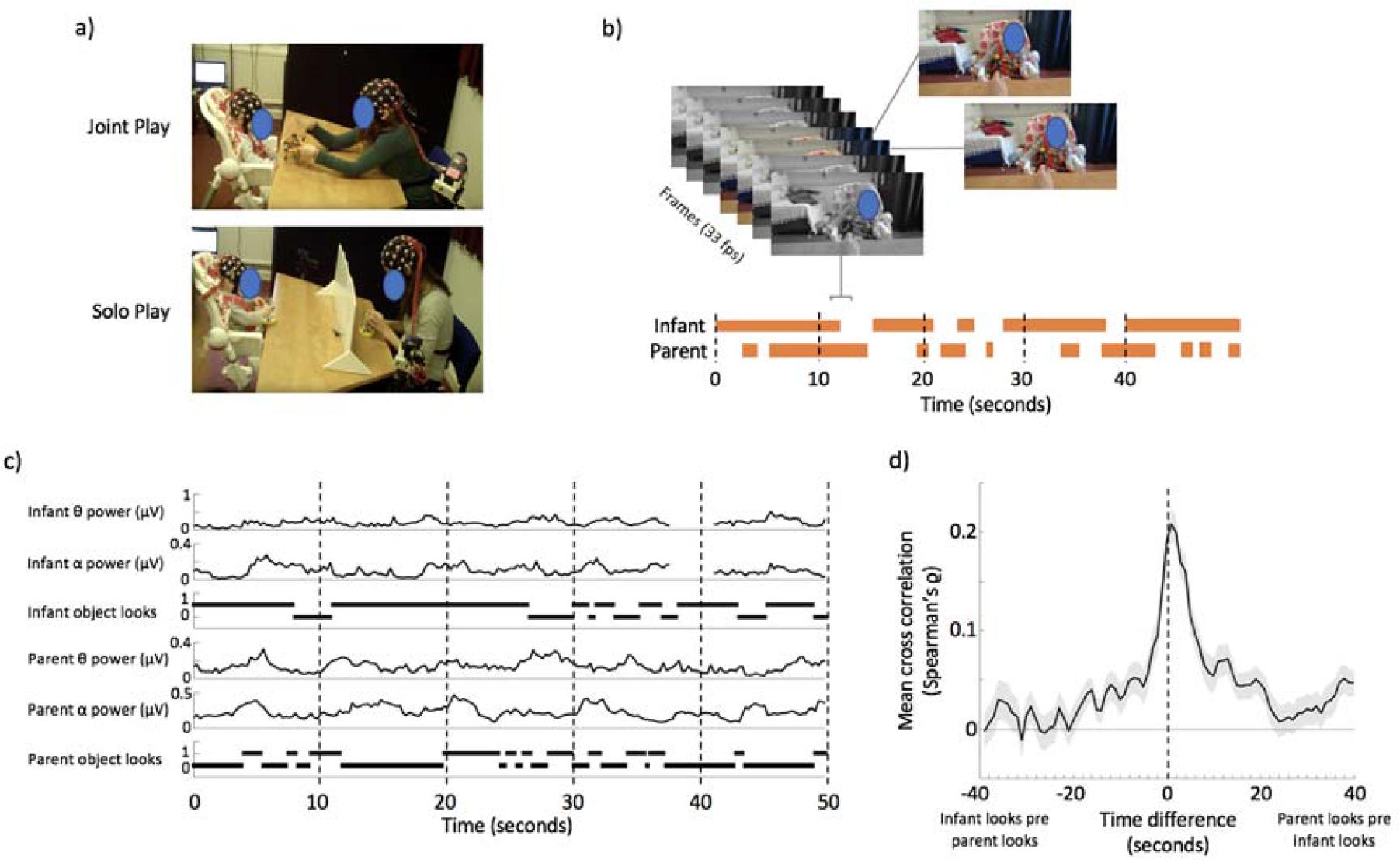
a) demonstration of experimental set-up; b) illustration of visual coding that was applied to the data; c) illustration of raw data. EEG data were decomposed using a Fourier decomposition and power within continuous bins was calculated, epoched to 4Hz; d) cross-correlation showing the relationship between infant object looks and parent object looks (see[25])

Based on previous research ([10, 13]) we expected that fluctuations in infant Theta activity would associate with, and forward-predict, fluctuations in infant attentiveness. Based on our previous research ([25]) we predicted that the forward-predictive relationship between infants’ own endogenous brain activity and infants’ attentiveness would be higher during Solo Play than Joint Play, due to the increased prevalence of exogenous parental attention scaffolding (and capture) during Joint Play. Further, since previous research indicates that parental responsiveness is an influential factor for early developing cognition ([35, 36]), we also examined whether parental neural responsivity had an effect on infants’ attention.

## Methods

*Participants*. 24 and 25 parents contributed usable data for the Joint Play (JP) and Solo Play (SP) conditions respectively; for infants, it was 21 and 25 for JP/SP respectively. Paired parent-child data were available for 20 dyads for Joint Play (10M/10F infants; mean (st.err.) infant age 345.1 (12.1) days; mother age 34.7 (0.8) years) and for 22 dyads for Solo Play (12M/10F infants; mean (st.err.) infant age 339.2 (10.3) days; mother age 34.1 (1.0) years). All participating parents were female. It should be noted that the recruitment area for this study, Cambridge, UK, is a wealthy university town and the participants were predominantly Caucasian and from well-educated backgrounds, and so do not represent an accurate demographic sample ([37]). Details of ethical permissions obtained are given in the Supplementary Materials.

*Experimental set-up*. Infants were seated in a high chair, and a table was positioned immediately in front so that toys on the table were within easy reach (see Figure 1). Parents were seated on the opposite side of the table, facing the infant. The table was 65cm wide. In the Solo Play condition only, a 40cm high barrier was positioned across the mid-line of the table. The height of the barrier ensured that, when it was in place, parent and child had line of sight to one another (to reduce the possibility of infant distress) but neither could see the others’ objects on the table.

Each infant-parent dyad participated in both the Joint Play and Solo Play conditions. Presentation order was randomised between participants, but the two conditions were always presented consecutively, with a short break between. Parents were informed that the aim of the study was to compare behaviour while they were attending to objects separately from each other, and when they were attending to the same object. During the Solo Play condition parents were asked to play silently with the toys separately from their infant, directing their own attention to the objects. During the Joint Play condition they were asked to play silently with the toys whilst involving their infant in the play,

A research assistant was positioned on the floor, out of the infant’s line of sight. The research assistant placed a series of toys onto the table, one at a time. In the Joint Play condition, one copy of each toy was presented to the infant and parent. In the Solo Play condition two identical toys were presented concurrently to the infant and parent, one on either side of the barrier. The toys were small (<15cm), engaging objects. The presentation order was randomised between conditions, and between participants. Approximately every two minutes, or more frequently if the child threw the object to the floor, the current toy object was replaced with a new object. The mean (st. err.) duration for which each object was presented was 140.1 (17.9) seconds for Joint Play and 110.3 seconds (7.9) for Solo Play. Approximately 10 minutes of data was collected per condition from each dyad. The mean (st.err.) duration of play for each condition was 10.80 (0.46) minutes for Joint Play and 10.35 (0.33) minutes for Solo Play. When the infant became fussy during testing, data collection was stopped earlier; however, this occurred fairly rarely: the number of infants contributing sessions that lasted less than 8 minutes was 2/3 for the Joint/Solo Play conditions.

*EEG Data Acquisition*. EEG signals were obtained using a 32-channel wireless Biopac Mobita Acquisition System and 32-channel Easycap. Further details of EEG acquisition are given in the Supplementary Materials.

*EEG Artifact rejection and pre-processing*. Automatic artifact rejection followed by manual cleaning using ICAs was performed. Full descriptions are given in the Supplementary Methods. Because previous analyses have shown that movement and muscle artefacts can contaminate EEG ([38, 39]), data from all channels other than the two channels close to the vertex, C3 and C4, were excluded and only frequencies between 2 and 14Hz were examined. Analyses suggested that these frequencies show least EEG signal distortion due to sweating, movement or muscle artefact ([38]). Prior literature (e.g. [11, 40]) suggests that these frequencies were also most likely to show associations with visual attention.

*EEG power analysis*. For each electrode, we computed the Fourier Transform of the activity averaged over artifact-free epochs, using the fast Fourier transform algorithm implemented in MATLAB (see Supplementary Materials for full description). The FFT was performed on data in 2000 ms epochs, which were segmented with an 87.5 % (1750 ms) overlap between adjacent epochs. Thus, power estimates of the EEG signal were obtained with a temporal resolution of 4 Hz and a frequency resolution of 1 Hz.

*Video coding and synchronisation*. Play sessions were videoed using two camcorders positioned next to the child and parent respectively. Further details of video coding and synchronisation are given in the Supplementary Materials. The visual attentional patterns of parents and infants was manually coded by reviewing their respective video recordings on a frame-by-frame basis (30 frames per second, 33.3 ms temporal acuity) using video editing software (Windows Movie Maker) (see Figure 1). This coding identified the exact start and end times of periods during which the participant was looking at the toy object.

*Calculation of time-lagged cross-correlation*. The attention data used for the cross-correlation analysis was re-sampled as continuous and time-synchronised data-streams at 4Hz (to match that of the EEG power estimate). Attention data were coded as 1 and 0 (either attentive towards the play object, or not). The cross-correlation calculations were performed separately for each frequency band (in 1Hz bands) and for each member of the dyad (infant brain-infant attention and parent brain-parent attention) (Analysis 1). Then, they were calculated across the dyad (parent brain-infant attention) (Analysis 2).

For each computation, the zero-lag correlation was first calculated across all pairs of time-locked (i.e. simultaneously occurring) epochs, comparing the EEG power profile with the attention data using a nonparametric (Spearman’s) correlation. (In the Supplementary Materials (Figure S3) we also show the results of the same tests repeated using alternative test, the Mann-Whitney U test, for which results were identical.) The mean correlation value obtained was plotted as time “0” (t=0) in the cross-correlation. Next, time-lagged cross-correlations were computed at all lags from −10 to +10 seconds in lags of +/-250ms (corresponding to one data point at 4 Hz). For example, at lag-time t=-250ms, the EEG power profile was shifted one data point backwards relative to the attention data, and the mean correlation between all lagged pairs of data was calculated. Based on an average of 10.5 minutes data per condition, sampled at 4Hz, and allowing for some attrition at artefact rejection due to the max-min thresholding criteria, the N of the cross-correlation was c.2300 for the zero-lag correlation and up to 40 fewer for the most shifted correlation. In this way, we estimated how the association between two variables changed with increasing time-lags. The individual cross-correlation series were then averaged across participants to obtain the group mean cross-correlation at each time interval and frequency band.

To compare the distribution of time x frequency data between any single condition and a null distribution, a cluster-based permutation test was conducted across time x frequency data, using the FieldTrip function ft_freqstatistics ([41]). In comparison to other approaches to solving the family-wise error rate, this approach identifies clusters of neighbouring responses in time/frequency space ([42]). In particular, corresponding time x frequency points were compared between contrast condition and null distribution with a t-test, and t values of adjacent spatiotemporal points with p<0.05 were clustered together with a weighted cluster mass statistic that combines cluster size and intensity (Hayasaka & Nichols, 2004). The largest obtained cluster was retained. Afterwards, the whole procedure, i.e., calculation of t values at each spatiotemporal point followed by clustering of adjacent t values, was repeated 1000 times, with recombination and randomized resampling before each repetition. This Monte Carlo method generated an estimate of the p value representing the statistical significance of the originally identified cluster compared to results obtained from a chance distribution.

In addition, a supplementary analysis was conducted using bootstrapping in order further to verify our results (see Supplementary Materials).

*Calculation of power changes around looks*. Analysis 3 examined whether individual looks accompanied by higher Theta power are longer lasting. To calculate this, we examined all looks to the play objects that occurred during the play session. The onset times of these looks were calculated, as described above, at 30Hz. Then, for each look, we excerpted the EEG power for three time windows immediately before, and after, the onset of each look (3000-2000, 2000-1000 and 1000-0 msecs pre look onset; 0-1000, 1000-2000 and 2000-3000 msec post look onset).

Separately, we calculated the duration of each look towards the object. Since these were heavily positively skewed, as is universal in looking time data ([43]), they were log-transformed. Then, we calculated separate linear mixed effects models for each of the six windows, using the *fitlme* function in Matlab. For each model we examined the relationship between EEG power within that time window and look duration, controlling for the random effect of participant. In this way we examined whether, for example, Theta power in the time window 1000 to 0 msec prior to the onset of a look showed a significant relationship to the subsequent duration of that look.

## Results

First, we examined the within-individual relationship between EEG power and visual attention, separately for Joint Play and Solo Play (Analysis 1). Second, to examine parental influences on infant attention, we calculated the cross-dyad cross-correlation between EEG power and visual attention (parent EEG power to infant attention), separately for Joint Play and Solo Play (Analysis 2). Third, we calculated changes in EEG power relative to individual look onsets (Analysis 3). Again, this was calculated both within-individual and across-dyad.

A previous report based on these data, that contained behavioural findings only, reported that infants showed longer look durations towards the object during Joint Play (JP) relative to Solo Play (SP), together with shorter periods of inattention (see Supplementary Figure S1) ([25]). Supplementary Figure S2 compares EEG power for infants and parents between Solo Play and Joint Play; no significant between-condition differences were observed.

### Analysis 1 – cross-correlation – within-participant

Figure 2 shows time-lagged cross-correlations between EEG power and visual attention for Solo Play. Figures 2a and 2b show correlations across the frequency spectrum, with time-lag on the x-axis and EEG frequency on the y-axis. Figures 2c and 2d show results of the cluster-based permutation test. These suggested that the results for both Infant Solo Play (p=.002) and Adult Solo Play (p=.002) differed significantly from chance. For infants, the effect was most pronounced in the 3-6 Hz range (Figure 2d); for adults, in the 6-12 Hz range (Figure 2e). In addition, to further confirm the results, a separate bootstrapping analysis was conducted, as described in the Supplementary Materials, which yielded identical results.

**Figure 2.**
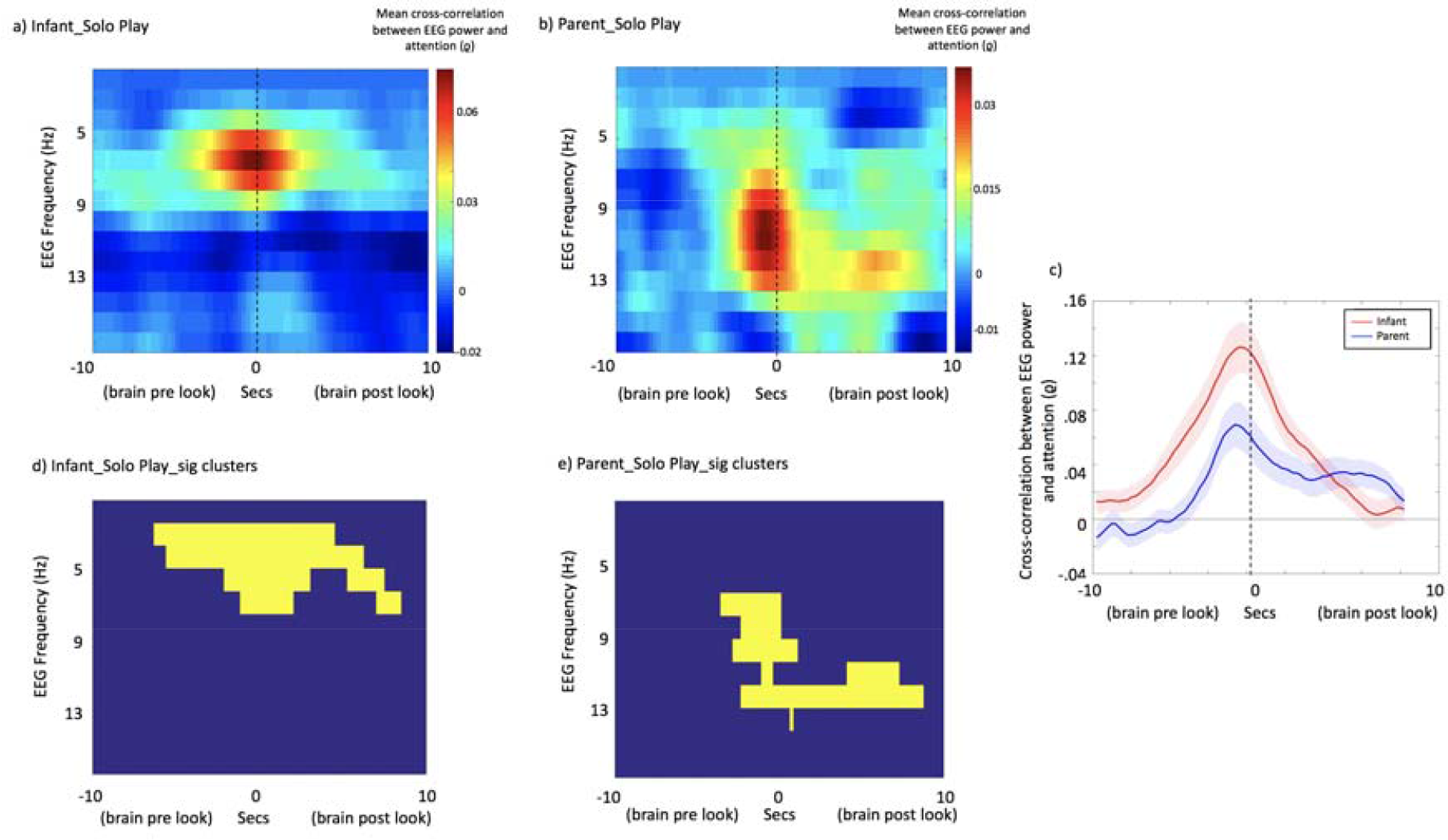
a and b – Mean time-lagged cross-correlations between EEG power and visual attention for a) Infant Solo Play and b) Parent Solo Play. Time lag between EEG power and visual attention is shown on the x axis and the EEG frequency on the y axis. c – Cross-correlation plots just for those frequency bands identified from the cluster-based permutation test as showing the most marked differences from chance (infant: 3-7Hz; adult: 6-12Hz). X-axis shows time; y-axis, cross-correlation between EEG power and attention. Shaded areas show the standard error of the means. d and e – results of the cluster-based permutation statistic. Yellow squares indicate time x frequency points of significant cross-correlations.

In order to examine at which time window the *peak* cross-correlation was observed between EEG power and visual attention, we excerpted the cross-correlation values just for those frequency bands identified from the cluster-based permutation test (infants: 3-7Hz; adults: 6-12Hz) (see Figure 2c). For infants, the peak cross-correlation was observed at t: −750ms (i.e. between EEG power at time t and attention 750 ms after time t). For adults, the peak cross-correlation was observed at t: −1000ms. (Of note, these numbers do not indicate the time lag of the EEG data relative to the *onset* of a look, but rather the time lag of the largest cross-correlation between EEG power and attention when treated as two continuous variables.)

Figure 3 compares the mean time-lagged cross-correlations for Infant Solo Play and Infant Joint Play. All data, including unpaired data, have been included (see Participants). Figures 3a and 3b show cross-correlation plots across the frequency spectrum. (Figure 3a is identical to 2a, and included to allow comparison with Figure 3b.) Figure 3d shows the cluster-based permutation test for the Infant Joint Play condition. This suggested that the Infant Joint Play condition differed significantly from chance (p=.008).

**Figure 3.**
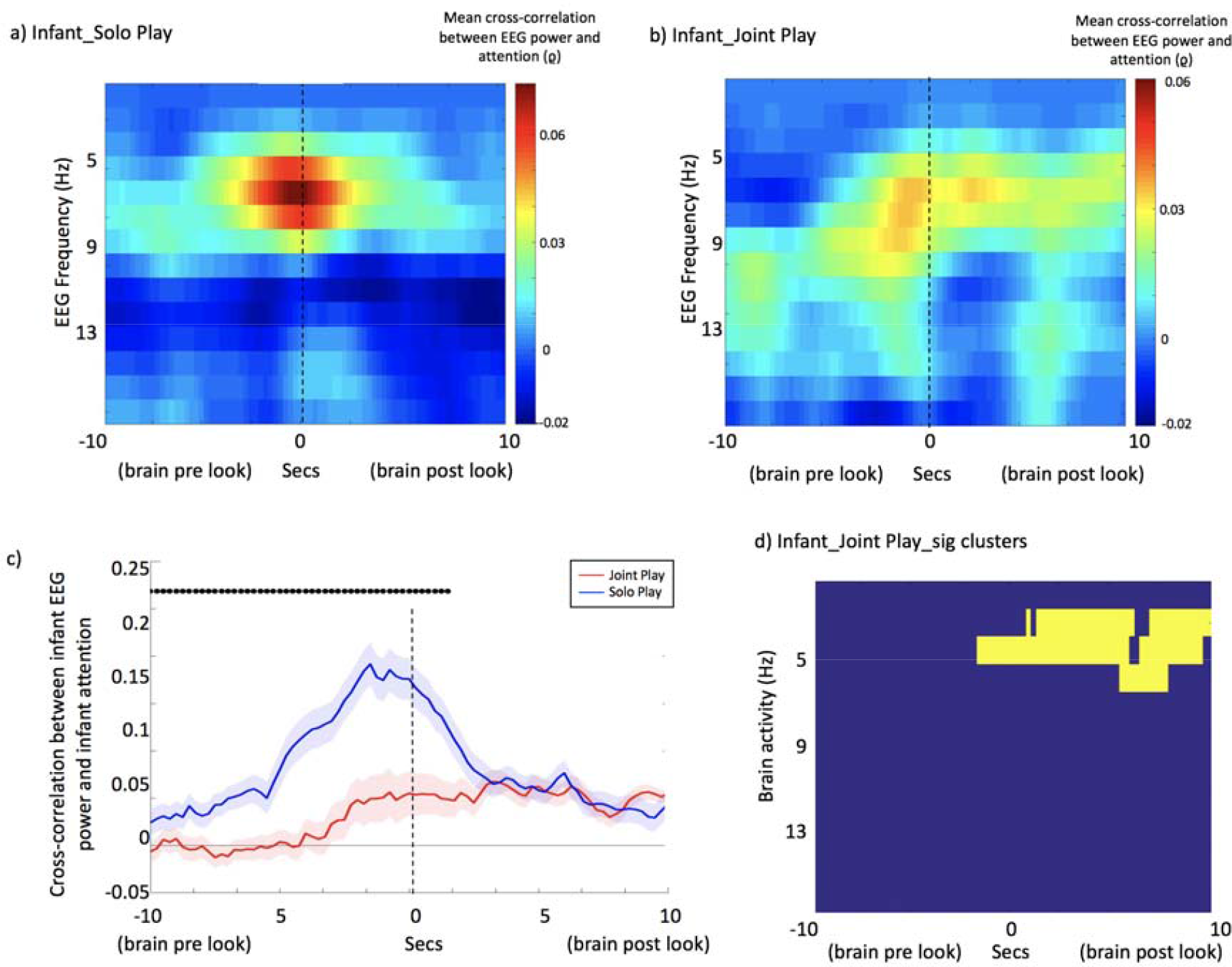
a and b – Mean time-lagged cross-correlations between EEG power and visual attention, for a) Infant Solo Play and b) Infant Joint Play. (Figure 3a is identical to Figure 2a, but included to allow for comparison with Figure 3b.) c - Line plot showing cross-correlation between EEG power and visual attention for just the frequency ranges identified from the cluster-based permutation test as showing marked effects in both conditions (3-6Hz). Red shows the Joint Play condition, and blue the Solo Play condition. Shaded areas show inter-participant variance (standard errors). Dots above the plots indicate the results of the significance calculations to assess whether the correlations observed differed significantly between the two conditions. d – results of the cluster-based permutation statistic for Infant Joint Play. Yellow squares indicate time x frequency points of significant cross-correlations.

To directly compare the *peak* cross-correlation values obtained for Infant Solo Play and Infant Joint Play, we excerpted the cross-correlation values just for those frequencies that the cluster-based permutation test indicated as showing marked differences in both conditions (3-6Hz) (see Figure 2c). For Solo Play, the peak cross-correlation was at t: −1500ms (EEG power at time t to attention 1500ms *after* time t); for Joint Play, the peak cross-correlation was at t: +3000ms.

In addition, separate unpaired t-tests were conducted at each time window to compare the results across conditions, and adjusted for multiple comparisons using the Benjamini-Hochberg FDR procedure ([44]). Time windows showing significant differences are indicated using black dots above the plot in Figure 3c. Results indicate that larger cross-correlations were observed during Solo Play relative to Joint Play for all time lags between t:-10,000ms and t: +1,250ms.

Figures 4a and 4b show the mean time-lagged cross-correlations for Parent Solo Play and Parent Joint Play. Figure 4e shows the cluster-based permutation test for Parent Joint Play, which indicated significant differences from chance (p=.001). For Parent Solo Play, the most marked associations between EEG power and attention were at 6-12 Hz (Figure 2b); for Parent Joint Play, the most marked associations were at 2-8 Hz (Figure 4e). To assess the significance of this difference we measured the frequency of peak association between EEG power and attention for parents during Solo Play and Joint Play, across all frequency bands under consideration (2-12Hz) during the +/- 1000msecs time window. Results obtained from the two conditions were compared using a paired t-test; a significant difference between the two conditions was observed t(44)=3.42, p=.001. This suggests that the peak association between brain activity and attention in the parent was observed at lower frequencies during Joint Play than during Solo Play.

**Figure 4:**
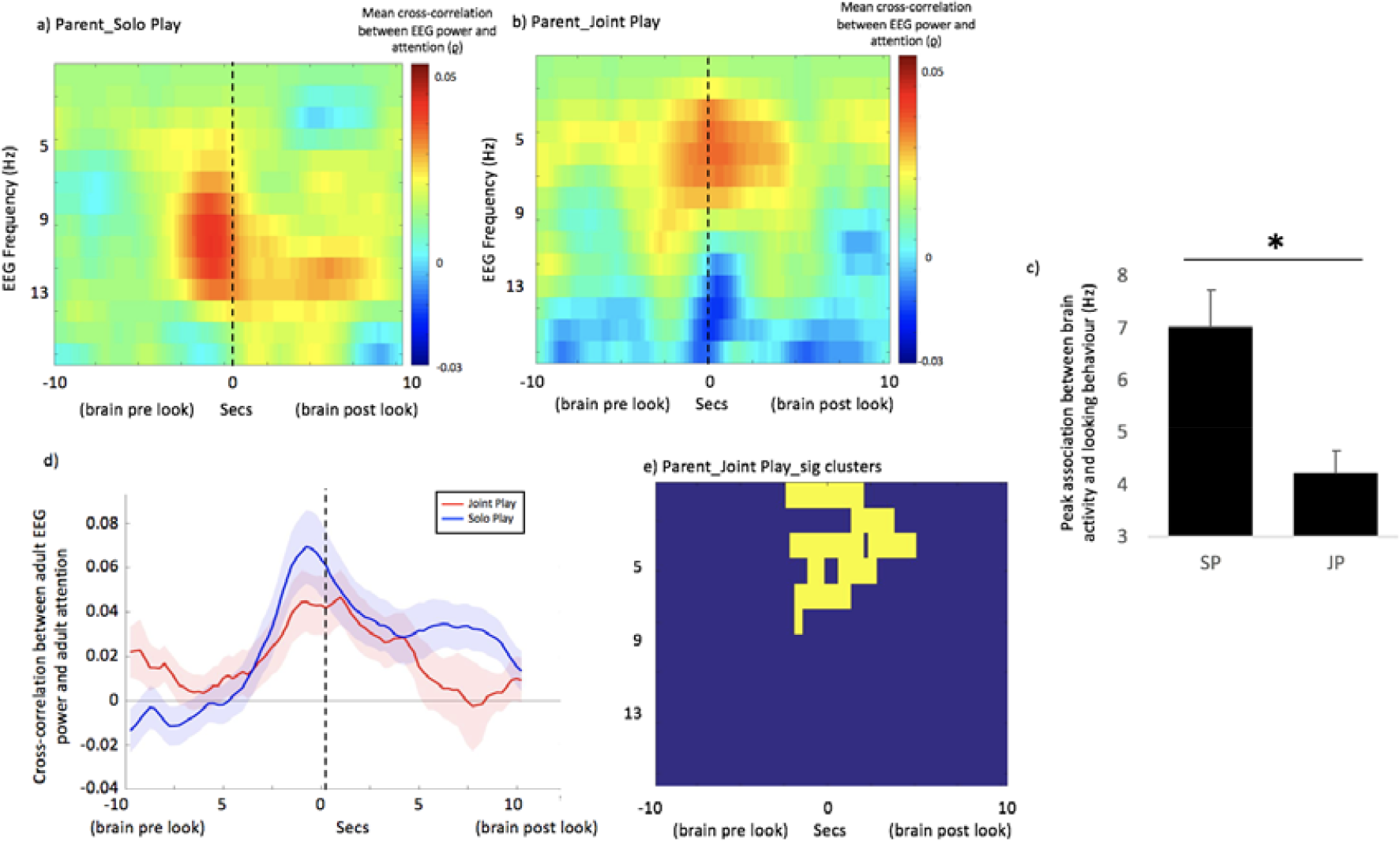
a and b – Mean time-lagged cross-correlations examining the relationship between EEG power and attention, for Parent Solo Play and Parent Joint Play. (Figure 4a is identical to Figure 2b, but scaled to be equivalent to Figure 4b to allow for comparison.) c - bar chart comparing the frequency of the peak association between EEG power and looking behaviour for parents in the Solo Play and Joint Play conditions. * indicates the results of the significance calculations, conducted as described in the main text. d - Line plot showing cross-correlation between EEG power and visual attention for just the frequency ranges identified from the cluster-based permutation test as showing marked effects in both conditions (Parent_Solo Play – 6-12Hz; Parent_Joint Play – 2-8Hz). Red shows the Joint Play condition, and blue the Solo Play condition. Shaded areas show inter-participant variance (standard errors). e - results of the cluster-based permutation statistic for Parent Joint Play. Yellow squares indicate time x frequency points of significant cross-correlations.

### Analysis 2 – cross-correlation – across parent and infant

Figures 5a and 5b show the mean time-lagged cross-correlations, and Figures 5d and 5e show the cluster-based permutation tests, for the relationship between parents’ EEG power and infants’ attention. For parent EEG and infant attention in the Joint Play condition a significant relationship was identified (p=.041). The most marked associations were identified in the 4-6Hz range (Figure 5e). An identical analysis examining the relationship between parent EEG and infant attention in the (concurrent but separate) Solo Play condition identified no significant relationship. In addition a further bootstrapping analysis was performed (see Supplementary Materials) which confirmed that the observed cross-correlation values significantly exceed chance for JP but not SP.

For the within-participant analysis of Solo Play, the peak-cross-correlation values observed were consistently *negative* (‘brain pre look’) (Figures 2c, 3c). In order directly to compare the *peak* cross-correlation values obtained between the Solo Play and Joint Play conditions, we excerpted the cross-correlation values just for those frequency bands identified from the cluster-based permutation test as showing marked differences during Joint Play (4-6Hz) (see Figure 5c). For Joint Play, the peak cross-correlation value occurred at a t:+750 ms (i.e. between infant attention at time t and adult EEG 750 ms *after* time t) (‘brain post look’).

**Figure 5:**
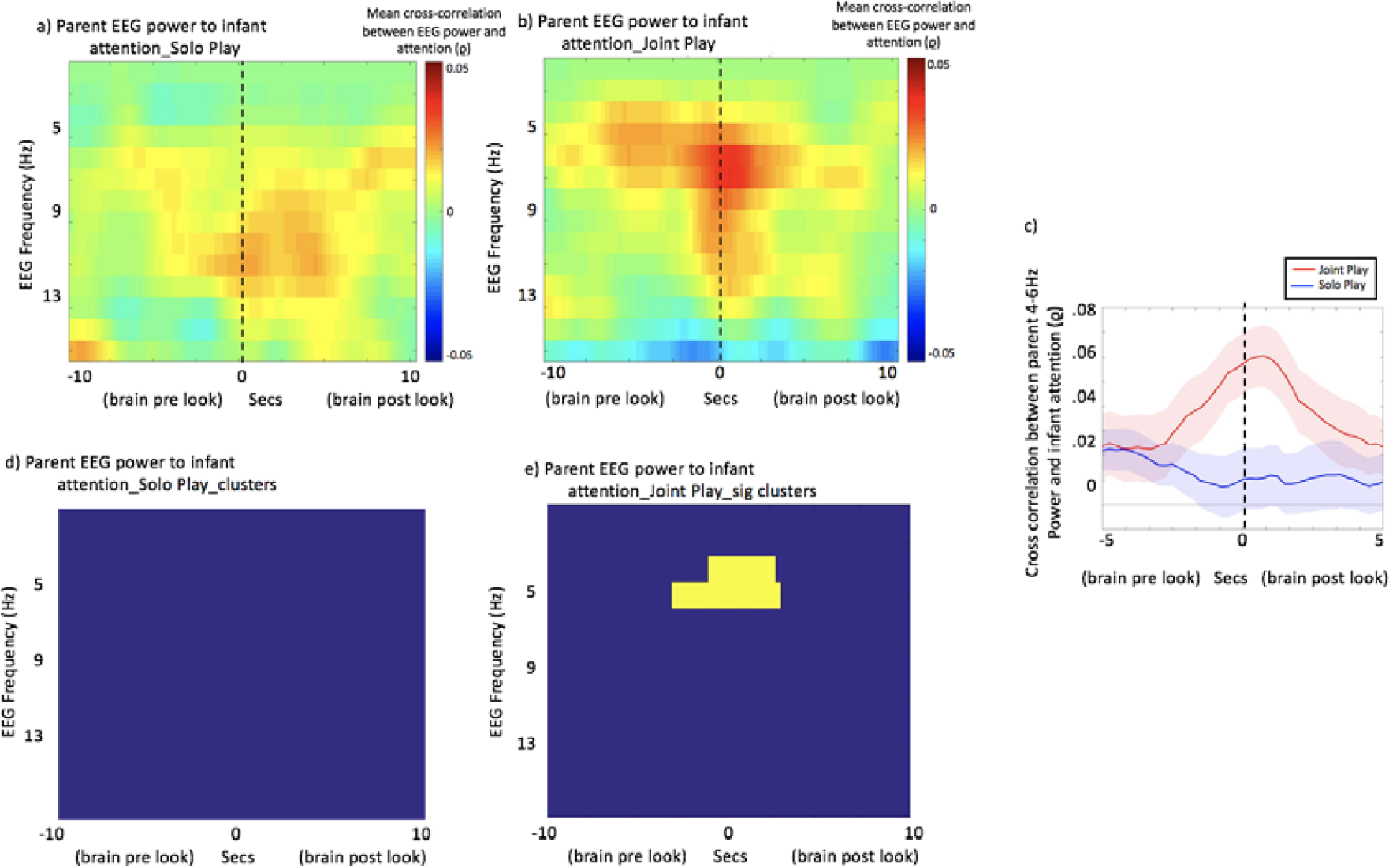
a and b – Mean time-lagged cross-correlations between parent EEG power and infant attention for a) Solo Play and b) Joint Play. Time lag between brain activity and visual attention is shown on the x axis and the EEG frequency on the y axis. c - Line plot showing cross-correlation between EEG power and visual attention for just the frequency ranges identified from the cluster-based permutation test as showing marked differences in the Joint Play condition (4-6Hz). Red shows the Joint Play condition, and blue the Solo Play condition. Shaded areas show inter-participant variance (standard errors). d and e – results of the cluster-based permutation statistic. Yellow squares indicate time x frequency points of significant cross-correlations.

### Analysis 3 – calculation of power changes around looks

In addition we conducted a further analysis using separate procedures to those used in Analyses 1 and 2. Whereas Analyses 1 and 2 examine the cross-correlation between EEG power and attention when treated as two continuous variables, analyses 3 examines changes in EEG power relative to the onsets of individual looks.

We examined all looks to the play objects that occurred during the session. For each look, we excerpted the power in the Theta band for the three time windows immediately prior to the onset of each look (3000-2000, 2000-1000 and 1000-0 msecs pre look onset) and the three windows immediately after the onset of each look (0-1000, 1000-2000 and 2000-3000 msec post look onset). Theta power was defined according to the frequency bands identified from the cluster-based permutation tests as showing the most marked differences from chance. These were: infant solo play (Figure 2d) - 3-7Hz; infant joint play (Figure 3d) - 4-7Hz); adult to infant (Figure 5e) - 4-6Hz.

We then calculated separate linear mixed effects models for each of the six windows, to examine the relationship between EEG power within that time window and look duration. Full results are shown in Supplementary Table S1, and key results are shown in Figure 6. In the Solo Play condition (Figure 6a) a relationship was observed between infants’ Theta power and look duration, consistent with the results of Analysis 1 (Figure 2a). Theta power in the time window −1000 to 0 msecs prior to look onset significantly predicted the subsequent duration of that look, consistent with the forward-predictive relationship noted in Figure 2c. The strength of this relationship increased for time windows after the onset of the look. Conversely, for Joint Play (Figure 6b), there was no significant relationship between infants’ Theta power and look duration. Again, this finding is consistent with the results of Analysis 1 (Figure 3c).

During Joint Play, parental Theta power associates significantly with infant attention in the time windows *after* the onset of the look (0 to 1000msec and 1000 to 2000msec) (Figure 6c) However there is no relationship in the time windows prior to look onset. This result is also consistent with the results of Analysis 2 (Figure 5c).

**Figure 6:**
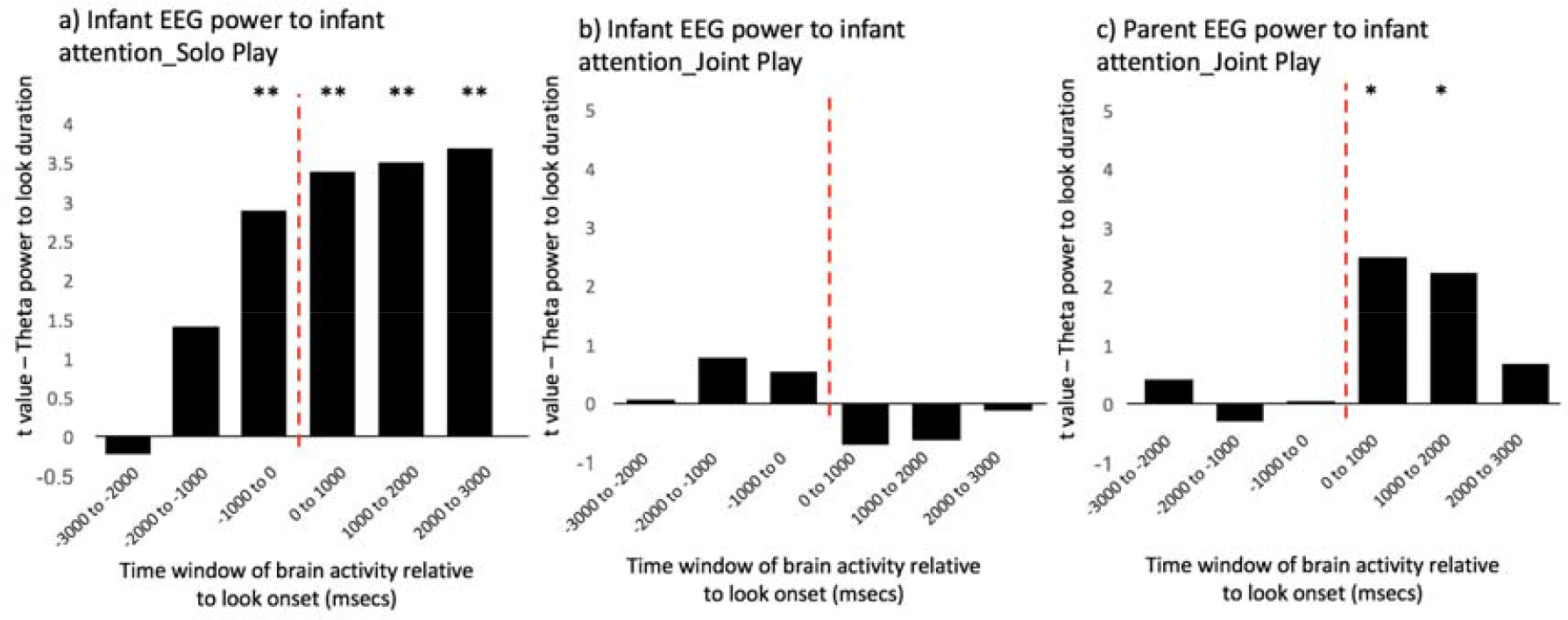
Results of linear mixed effects models conducted to examine whether individual looks accompanied by higher Theta power are longer lasting. For each look, the Theta power for three time windows prior to look onset (3000-2000, 2000-1000 and 1000-0 msecs pre look) and for three time windows post look onset (0-1000, 1000-2000 and 2000-3000 msec post look) was excerpted. We then calculated separate linear mixed effects models for each of the six windows, to examine the relationship between EEG power within that time window and look duration. Y axis shows the t value. *indicates the p values (*<.05, **<.01). Full results are shown in Supplementary Table S1.

## Discussion

It is well established that attention and learning are supported by the endogenous oscillatory neural activity of the person attending. However, relative little is known about how interpersonal and social influences on attention are substantiated in the brain ([16, 45]). To investigate this, we examined how the oscillatory dynamics of attention are shared between infant-parent dyads, and how these dynamics differ between non-interactive and interactive social play.

We found that, when infants were engaged in Solo Play, continuous fluctuations in Theta power forward-predicted visual attention in infants (Figure 2). Consistent with this, a separate analysis identified a positive association between Theta power in the 1000ms prior to look onset and the subsequent duration of that look (Figure 6). For adults, a similar functional relationship was observed, but at a higher frequency (6-12 Hz) in the Alpha band, consistent with considerable previous research into the role of pre-stimulus Alpha activity in anticipatory visual attention ([46, 47]). Our infant findings are also consistent with previous research suggesting that Theta oscillations increase in during anticipatory and sustained attention ([10]; [12, 13]); but they are novel insofar as we demonstrated these effects during spontaneous attention in semi-naturalistic settings.

During interactive, social play, however, we found that this forwards-predictive relationship between infants’ endogenous Theta activity and visual attention was still present, but much reduced. Again, this result was observed consistently across two separate analyses (Figure 3 and Figure 6). Particularly of interest was Figure 3c, which suggested that negative-lag relationships (attention forward-predicting EEG power) were similar across the Solo and Joint Play conditions, but that positive-lag relationships (EEG power forward-predicting attention) were present only during Solo Play. These results are consistent with our previous research suggesting that endogenous factors, such as attentional inertia, influence infants’ attention more during solo (non-interactive) play than during joint play ([25]). Taken together, our results suggest that infants’ endogenous neural control over attention is greater during solo play.

These results appear unlikely to be attributable to oculomotor artefact associated with the onsets and offsets of looks, for a number of reasons. First, during data pre-processing we removed oculomotor artifacts via ICA (see Supplementary Materials); second, we have only reported data in this paper from two channels near the vertex – C3 and C4, which show the least contamination by muscle and motion artifacts. Third, the cross-correlation analysis across different frequencies (Figure 2a) indicated that relationships were specific to the Theta band. Muscular artefact generally produces the highest contamination in Delta, Beta and Gamma bands ([38, 39]). Fourth, effects were present around the onsets of looks in the Solo Play, but not the Joint Play, condition (Figure 3a, 3b).

Our findings are also unlikely to be attributable to differences in mean look duration between the two conditions (see Figure S1), for two reasons. First, as in Analysis 1, any artifactual effects would be random rather than directional (i.e. specifically affecting negative rather than positive lags). Second, Analysis 1 examined the relationship between attention and EEG power considered across continuous entire time series, whereas Analysis 3 examined power changes relative to the onsets of individual looks, and the results from the two analyses produced converging conclusions. Furthermore, this result is also not attributable to differences in relative power between the two conditions, as the EEG power spectrum of infants did not differ across conditions (Figure S2).

Overall, however, we found that, despite the fact that infants’ endogenous attention control over their own behaviour patterns appeared to be lower, they were *more attentive* towards objects during Joint Play – a finding consistent with previous research ([24]). To understand why, we examined how adult brain activity related to infant attention.

First, we found that, during Joint Play, the frequency of adults’ peak association between EEG power and attention was down-shifted to the Theta range – similar to infants’ peak frequency of association (Figure 4). Second, we found that parent EEG Theta power significantly co-fluctuated with infant attention. Again, this result was observed across two separate analyses. Analysis 2 (Figure 5d, 5e) suggested that infant attention associated, over a time-frame of +/− 2 seconds, with increased parental Theta power. Analysis 3 (Figure 6c) suggested that individual infant attention episodes accompanied by greater parental EEG power were longer lasting.

Importantly, we found that the direction of the peak association differed between solo and interactive play. During Solo Play, the peak cross-correlation between infant Theta power and infant attention was observed at negative lag (‘brain pre look’) (Figure 2c, 3c), and Theta power 1000ms prior to look onset predicted look durations (Figure 6c). During Joint Play, the peak cross-correlation between adult Theta power and infant attention was observed at positive lag (‘brain post look’) (Figure 5c), and Analysis 3 identified backwards-predictive but not forwards-predictive relationships between adult Theta power and infant look duration (Figure 6c). These findings appear to suggest that, during Joint Play, parents’ Theta power tracks, and responds to, changes in infants’ attention.

One possible account of our findings we considered is that infant attention may (Granger-) cause adult attention, which in turn causes increased Theta activity in adults. This explanation appears unlikely however, because in the Supplementary Materials we report a control analysis where instances in which an attention shift from the infant was immediately followed by an attention shift from the parent were excluded. The results obtained from this subset of the data were highly similar to those reported in the main text (see Figure S5). Furthermore, as we show in Figure 1d, adults’ gaze forward-predicted infants’ attention more than *vice versa*, which also appears inconsistent with this explanation.

Overall, then, our results suggest that adults show neural responsivity to the behaviours of the child, and that increased parental neural responsivity associates, look-by-look, with increased infant attentiveness. Temporally fine-grained patterns of parental responsivity to infants have previously been shown using methods other than neuroimaging, such as micro-coding of facial affect ([48, 49]), autonomic physiology ([50]), visual attention ([51]) and vocalisations ([52]; [53]). And, using neuro-imaging, research with adults has provided evidence for common activation elicited when experiencing emotions such as disgust ([54]), touch ([55]) or pain ([56]) in oneself, and when perceiving the same feelings in others. However, this is the first study, to our knowledge, to demonstrate temporal associations between infants’ attentiveness and parental neural correlates of attention, and to show that moment-to-moment variability in adults’ neural activity associates with moment-to-moment variability in infants’ attentiveness.

Although demonstrated here in the context of parent-child interaction, future research should explore whether our present findings extend to cover other aspects of social interaction ([57]). They should also be extended to explore individual differences – whether some social partners show greater neural responsiveness to others, and how this influences behaviour ([49]) – and to other aspects of inter-personal neural influences than shared attention during joint play. Finally, future work should examine the mechanisms through which the children of parents who show increased responsivity over shorter time-frames develop superior endogenous attention control over long time-frames ([21–23, 58, 59]).

## Author contributions

SW and VL designed research. SG, KC, LB, RN, VL, VN, LS performed research. SW and VN performed analyses. SW and VL wrote manuscript.

## Declaration of conflicting interests

None declared.

## Acknowledgements

This research was funded by ESRC Grant numbers ES/N006461/1 to VL and SW, ES/N017560/1 to SW, and Nanyang Technological University Grant M4081585.SS0 to VL. Thanks to John Duncan and Paul Chadderton for commenting on early versions of this manuscript.

